# CAGEcleaner: reducing genomic redundancy in gene cluster mining

**DOI:** 10.1101/2025.02.19.639057

**Authors:** Lucas De Vrieze, Miguel Biltjes, Sofya Lukashevich, Kodai Tsurumi, Joleen Masschelein

## Abstract

**Summary:** Mining homologous biosynthetic gene clusters (BGCs) typically involves searching colocalised genes against large genomic databases. However, the high degree of genomic redundancy in these databases often propagates into the resulting hit sets, complicating downstream analyses and visualisation. To address this challenge, we present CAGEcleaner, a Python-based tool with auxiliary bash scripts designed to reduce redundancy in gene cluster hit sets by dereplicating the genomes that host these hits. CAGEcleaner integrates seamlessly with widely used gene cluster mining tools, such as cblaster and CAGECAT, enabling efficient filtering and streamlining BGC discovery workflows.

**Availability and implementation:** Source code and documentation is available at GitHub (https://github.com/LucoDevro/CAGEcleaner) and at Zenodo (https://doi.org/10.5281/zenodo.14726119) under an MIT license. CAGEcleaner comes with its own Conda environment but can also be installed from the Python Package Index (https://pypi.org/project/cagecleaner/).

**Supplementary information:** Supplementary data are available at *Bioinformatics* online.

## Introduction

Biosynthetic pathways are often driven by multiple genes that are physically grouped together in the genome. Across all kingdoms of life, such gene clusters are extensively studied for their ability to direct the biosynthesis of diverse metabolites and proteins in a streamlined fashion. They play a central role in various biological processes, such as secondary metabolism, virulence, toxin production and drug resistance. Comparative analysis of gene clusters provides valuable insights into their evolutionary trajectories, and the functional and biosynthetic diversity of the pathways they encode. Tools such as MultiGeneBlast (Medema, Takano and Breitling 2013), antiSMASH (Blin et al. 2023) and, most recently, cblaster (Gilchrist et al. 2021) and CAGECAT (van den Belt et al. 2023), have facilitated such large-scale comparative analyses. These tools can provide a wide view on gene cluster diversity by querying large public genome databases, such as those hosted by NCBI. However, these large databases contain substantial genomic redundancy due to the deposition of (re)sequenced (nearly) identical genomes, as well as from continuous sequencing efforts in the context of pathogen surveillance. This redundancy tends to propagate into the output of gene cluster mining tools, often yielding hundreds of quasi-identical clusters. As a result, a tedious curation process is typically required before meaningful downstream analyses and visualisations can be performed.

Here, we present CAGEcleaner, a Python-based pipeline with auxiliary bash scripts that rapidly dereplicates gene cluster sets by assessing the genomic similarity of their host genomes. In addition, it can preserve a certain degree of genomic redundancy when justified by sufficient gene cluster diversity. Designed primarily as a post-processing tool for cblaster, CAGEcleaner serves as an intermediate filtering step, streamlining downstream analyses and visualisations, including those facilitated by cblaster’s sister package clinker (Gilchrist and Chooi 2021).

### Implementation

The CAGEcleaner workflow is outlined in Figure 1 and consists of three parts. As input, CAGEcleaner requires a cblaster session file in json-format, obtained after running a search query.

**Figure 1:**
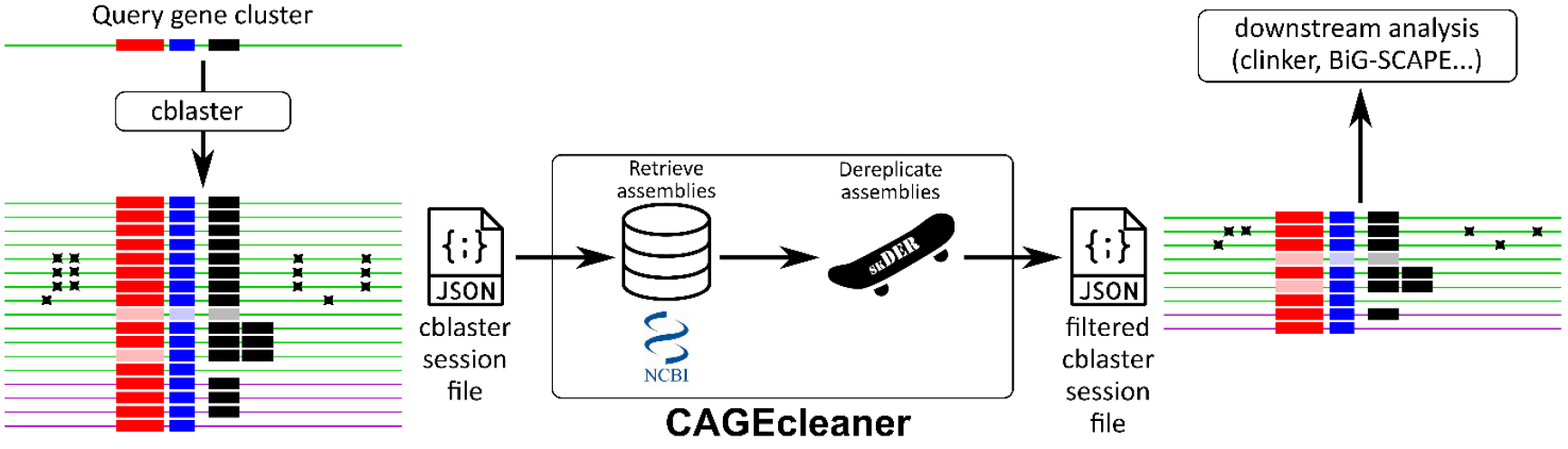
Schematic overview of the CAGEcleaner pipeline. A cblaster search query of three colocalised genes (red, blue and black rectangles) returned several redundant cluster hits. Some of these are from the same species as the query (green lines), while other hits are from another species (purple lines). Some of the same-species hits are hosted by strains that are only remotely related to the query strain, which, for example, shows in the presence of point mutations (black crosses). Starting from the session file of this search, CAGEcleaner dereplicates the genomes that host these hits, and returns a filtered session file that can readily be used for downstream analyses and visualisations. It preserves some degree of genomic redundancy when justified by (1) outlier cblaster homology scores (hits with transparent rectangles), or (2) a different number of query gene homologs than in the query cluster (hits with two or none black rectangles).

In the first part, the genome assemblies associated with the cblaster hits are retrieved. The scaffold NCBI Nucleotide IDs are first extracted from the cblaster session file and mapped to NCBI Assembly IDs using the Entrez-Direct utilities, executed via an auxiliary bash script. Scaffold entries that are part of an NCBI Whole Genome Shotgun (WGS) project are first redirected to their respective WGS master records before being mapped to Assembly IDs. The mapped assemblies are then downloaded as gzipped nucleotide FASTA files using the NCBI Datasets CLI (O’Leary et al. 2024) by another auxiliary bash script. To speed up ID mapping, we make the Entrez-Direct utilities process large batches of 5000 scaffold IDs at once. However, the Entrez-Direct utilities do not preserve the mapping between scaffold IDs and assembly IDs when mapping in batches. Since maintaining this mapping is critical for pinpointing scaffolds hosting gene cluster hits retained after genome dereplication, CAGEcleaner reconstructs this mapping locally by matching each scaffold ID extracted from the cblaster session file to a scaffold ID contained within one of the retrieved assembly FASTA files.

The second part of the workflow involves the dereplication of the genome assemblies. This process is performed using skDER, a fast genome dereplication tool which clusters genome assemblies based on pairwise average nucleotide identity (ANI), aligned fraction (AF), assembly N50 and connectedness metrics, and selects a representative genome for each genome cluster (Salamzade and Kalan 2023). In an auxiliary bash script, skDER is run in greedy mode on the locally downloaded genome assemblies using a user-defined ANI threshold. The resulting output tables are then parsed to identify the skDER representative genome assemblies.

The third part of the workflow identifies the gene cluster hits to be retained by mapping the IDs of the representative genome assemblies back to gene cluster scaffold IDs using the earlier reconstructed scaffold-assembly ID mapping. In addition, it recovers gene clusters within redundant, non-representative genome assemblies that exhibit sufficient cluster diversity to justify their retention in the final gene cluster hit set. This ensures the preservation of gene cluster diversity within highly similar genomes, which may have arisen through recent genomic reorganisation or horizontal gene transfer. Such hits are detected using two strategies. The first strategy evaluates the contents of each gene cluster by subdividing each skDER-identified genome cluster into subgroups based on the number of homologs for each gene in the query cluster. A new representative genome is then randomly selected from each subgroup and retained in the hit set, skipping the subgroup including the earlier retained skDER representative. The second strategy assesses the cblaster homology scores of each gene cluster hit. Within each subgroup, hits with outlier scores as identified using z-scoring, are retained, ensuring that functionally distinct gene clusters are not removed.

Finally, CAGEcleaner generates seven output text files summarising the dereplicated gene cluster hit sets. Intermediate output, such as downloaded genomes or skDER results, can also be returned upon request. These seven output text files are described in Table 1.

**Table 1:**
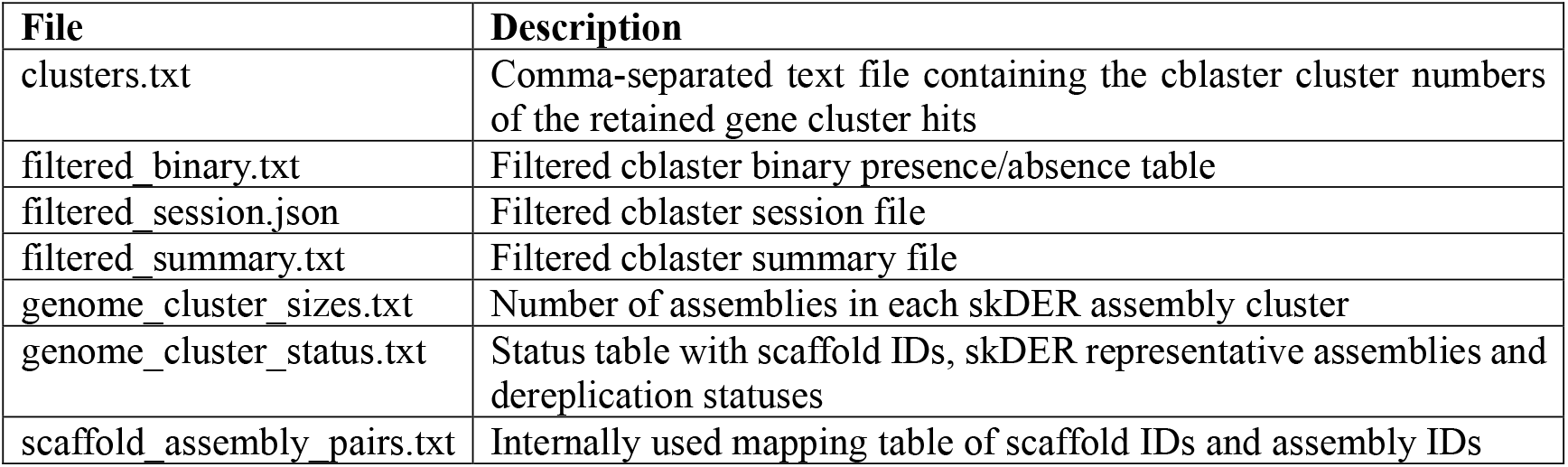
Description of the seven CAGEcleaner output text files.

| File | Description |
| --- | --- |
| clusters.txt | Comma-separated text file containing the cblaster cluster numbers of the retained gene cluster hits |
| filtered_binary.txt | Filtered cblaster binary presence/absence table |
| filtered_session.json | Filtered cblaster session file |
| filtered_summary.txt | Filtered cblaster summary file |
| genome_cluster_sizes.txt | Number of assemblies in each skDER assembly cluster |
| genome_cluster_status.txt | Status table with scaffold IDs, skDER representative assemblies and dereplication statuses |
| scaffold_assembly_pairs.txt | Internally used mapping table of scaffold IDs and assembly IDs |

### Example cases

We evaluated the running time, disk usage and RAM usage of CAGEcleaner for two concrete example cases. In each case, we executed the workflow using 20 CPU cores (Intel Core i7-13700k) and provided 32 GB of RAM.

In case 1, we queried four genes - the two core biosynthetic genes and their direct neighbours - from the actinorhodin biosynthetic gene cluster from *Streptomyces coelicolor A3(2)* (MIBiG (Terlouw et al. 2023) entry BGC0000194) against the NCBI RefSeq Protein database using cblaster at default settings. This yielded 8934 gene cluster hits in the binary table. After running CAGEcleaner at its default settings (ANI threshold of 99%), this hit set was reduced to 4847 hits, representing a 1.84-fold reduction. Among the retained hits, 170 were recovered by cluster content and 11 by outlier score. This run required 1 hour 29 minutes, consuming 28.5 GB of disk space and 27.6 GB of memory. In case 2, we aimed to have more redundancy in the hit set. We performed a generic query of three colocalised *Staphylococcus* genes against NCBI RefSeq Protein using cblaster at default settings and applied an Entrez query filter “Staphylococcus[orgn]”. This yielded 1146 gene cluster hits., which CAGEcleaner reduced to just 22 hits, a 52-fold reduction. The run required 10 minutes, 1.2 GB of disk space and 1.7 GB of memory.

The inputs and outputs for these two example cases are provided as Supplementary Material.

## Discussion

Hit redundancy is a persisting challenge in genome mining, often complicating visualisation and downstream analyses. CAGEcleaner is the first tool capable of tackling this issue in an automated manner. By leveraging efficient genome dereplication strategies, it significantly reduces large gene cluster hit sets in a limited amount of time. CAGEcleaner has been deliberately designed to dereplicate on the host genome level, rather than on the gene cluster level. As such, it can discern different host species and/or strains, and preserve the genomic evolutionary signal throughout the dereplication process, opening up new avenues for downstream analysis. For example, by contrasting genome-level clustering from skDER with the clustering output from gene cluster-level analysis tools like BiG-SCAPE (Navarro-Muñoz et al. 2020), it may provide insights into horizontal gene transfer events, or uncover gene clusters that evolve at different rates compared to the overall genome. CAGEcleaner is implemented in Python 3 and bash, and can be freely installed from GitHub and the Python Package Index https://pypi.org/project/cagecleaner/). Source code and documentation is available on the GitHub page (https://github.com/LucoDevro/CAGEcleaner).

## Supporting information

Actinorhodin Example Case Files

Staphylococcus Example Case Files

## Acknowledgement

The authors thank Cameron Gilchrist for his helpful advice on seamlessly integrating CAGEcleaner with cblaster.

